# *In vivo* structures of an intact type VI secretion system revealed by electron cryotomography

**DOI:** 10.1101/108233

**Authors:** Yi-Wei Chang, Lee A. Rettberg, Grant J. Jensen

## Abstract

The type VI secretion system (T6SS) is a versatile molecular weapon used by many bacteria against eukaryotic hosts or prokaryotic competitors. It consists of a cytoplasmic bacteriophage tail-like structure anchored in the bacterial cell envelope via a cytoplasmic baseplate and a periplasmic membrane complex. Rapid contraction of the sheath in the bacteriophage tail-like structure propels an inner tube/spike complex through the target cell envelope to deliver effectors. While structures of purified contracted sheath and purified membrane complex have been solved, because sheaths contract upon cell lysis and purification, no structure is available for the extended sheath. Structural information about the baseplate is also lacking. Here we use electron cryotomography to directly visualize intact T6SS structures inside *Myxococcus xanthus* cells. Using sub-tomogram averaging, we resolve the structure of the extended sheath and membrane-associated components including the baseplate. Moreover, we identify novel extracellular bacteriophage tail fiber-like antennae. These results provide new structural insights into how the extended sheath prevents premature disassembly and how this sophisticated machine may recognize targets.

## MAIN TEXT

The type VI secretion system (T6SS) is a dynamic nanomachine, widespread in Gram-negative bacteria, that delivers effectors directly into target eukaryotic or bacterial cells for purposes of pathogenicity or competition^1-3^. Its structure is known to comprise a membrane complex linked to the outer membrane (OM) and spanning the periplasm and inner membrane (IM), a cytoplasmic baseplate associated with the membrane complex, an extended sheath anchored to the baseplate and loaded with an inner tube made of stacks of hexameric Hcp rings, and a spike complex on the tip of the inner tube^1-3^. The extended sheath contracts to fire the inner tube/spike complex into target cells to deliver associated effectors^4-6^.Bioinformatics and structural studies have provided strong evidence that this contractile mechanism is conserved with the contractile bacteriophage tail, R-type pyocin, and phage tail-like protein translocation structures^7^.

Among these contractile machines, the T6SS is the only one that fires from inside the cell, and the only one known to recycle its components for multiple rounds of action^4,8^. The sheath proteins of the T6SS are equipped with a unique recycling domain recognized by the ATPase ClpV for rapid disassembly of the contracted sheath, allowing the sheath subunits to be reused to build other extended T6SS^6,9,10^. High-resolution cryo-EM reconstructions have been obtained for purified R-type pyocins in both the extended and contracted conformations, revealing how the energy for contraction is stored in the extended sheath and released upon contraction, and how the extended sheath interacts with the inner tube^11^. For the T6SS, however, extended sheaths have not been reconstructed because cell lysis and purification trigger contraction. As a result, only the structure of the contracted sheath has been obtained by high-resolution cryo-EM^12,13^. Consequently, how the T6SS sheath subunits pack in the extended state, manage their recycling domains to prevent premature disassembly by ClpV, and interact with the Hcp inner tube are still unknown. In addition, the structural details of T6SS attachment to the membrane remain unclear.

Electron cryotomography (ECT) has proven powerful in visualizing native structures of bacterial nanomachines directly in their cellular context without purification^14^. In fact, the structure of the T6SS was first discovered by ECT in *Vibrio cholerae* cells, wherein both extended and contracted conformations were seen, revealing the basic contractile mechanism^4^. Following this discovery, we used correlated cryogenic photoactivated localization microscopy and ECT (cryo-PALM/ECT) to identify the T6SS structure *in vivo* in *Myxococcus xanthus* ^15^, which was known to encode a T6SS in its genome Fig. S1). By fusing photoactivatable GFP with the sheath, localizing the signals in a frozen-hydrated cell, and correlating the signals to electron densities in a cryotomogram of the same cell, we identified T6SS and confirmed that they formed extended and contracted structures just like their V. cholerae counterparts. We further noticed that in data taken close to focus, clear periodicities were visible in the M. xanthus T6SS sheaths (Fig. S2), opening the possibility of sub-tomogram averaging to reveal structural details.

**Figure 1.**
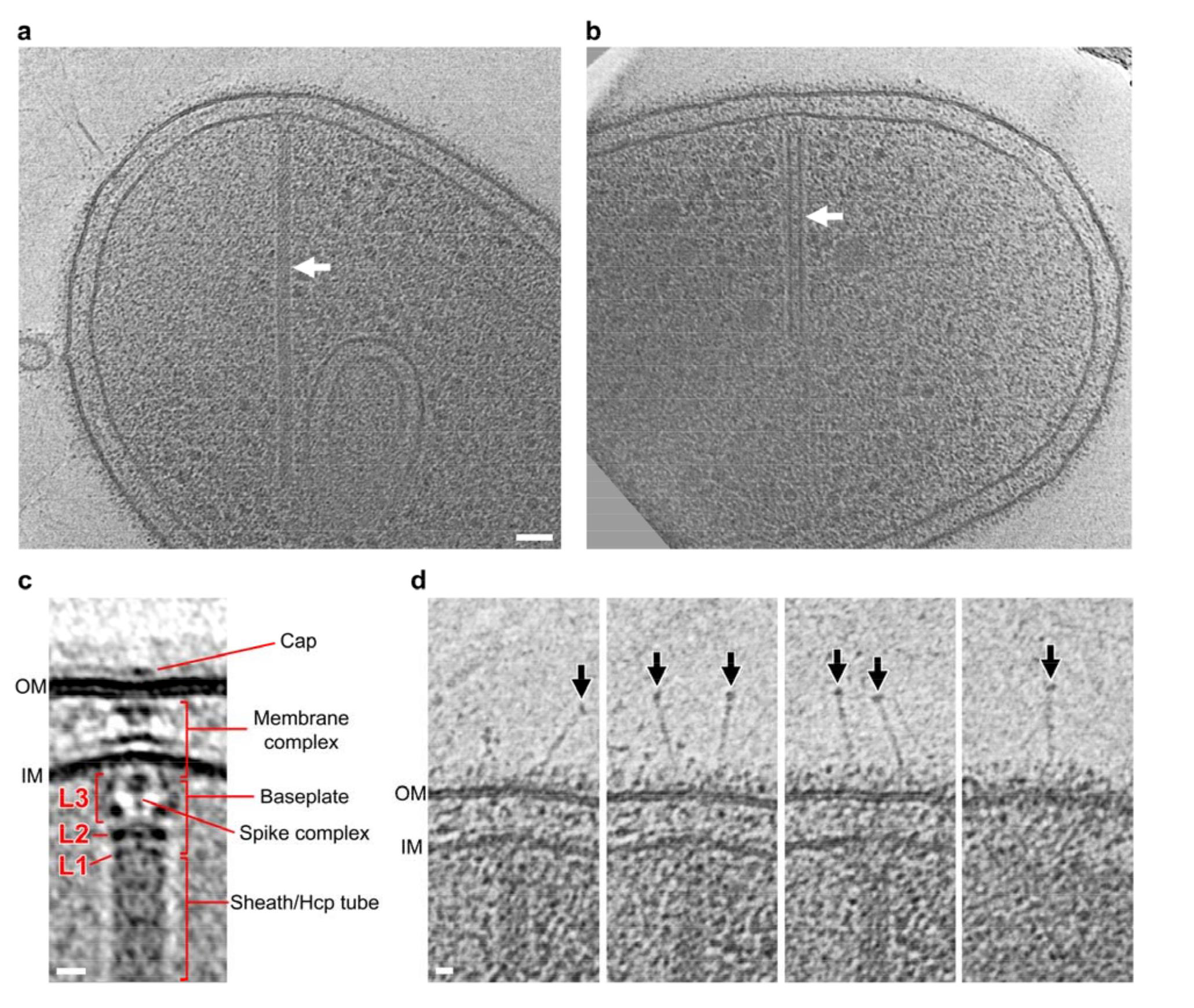
Visualization of the *M. xanthus* T6SS *in vivo*. (a, b) Tomographic slices through an extended (a) and a contracted (b) T6SS (arrows) in frozen376 hydrated M. xanthus cells. (c) Central slice of the sub-tomogram average of the extended T6SS.L1-L3: different layers of densities in between the sheath and the IM. (d) Extracellular bacteriophage tail fiber-like antennae captured in different slices of the tomogram shown in (a).Scale bar in (a) 50 nm, applies to (a) and (b); scale bars in (c) and (d) 10 nm.

**Figure 2.**
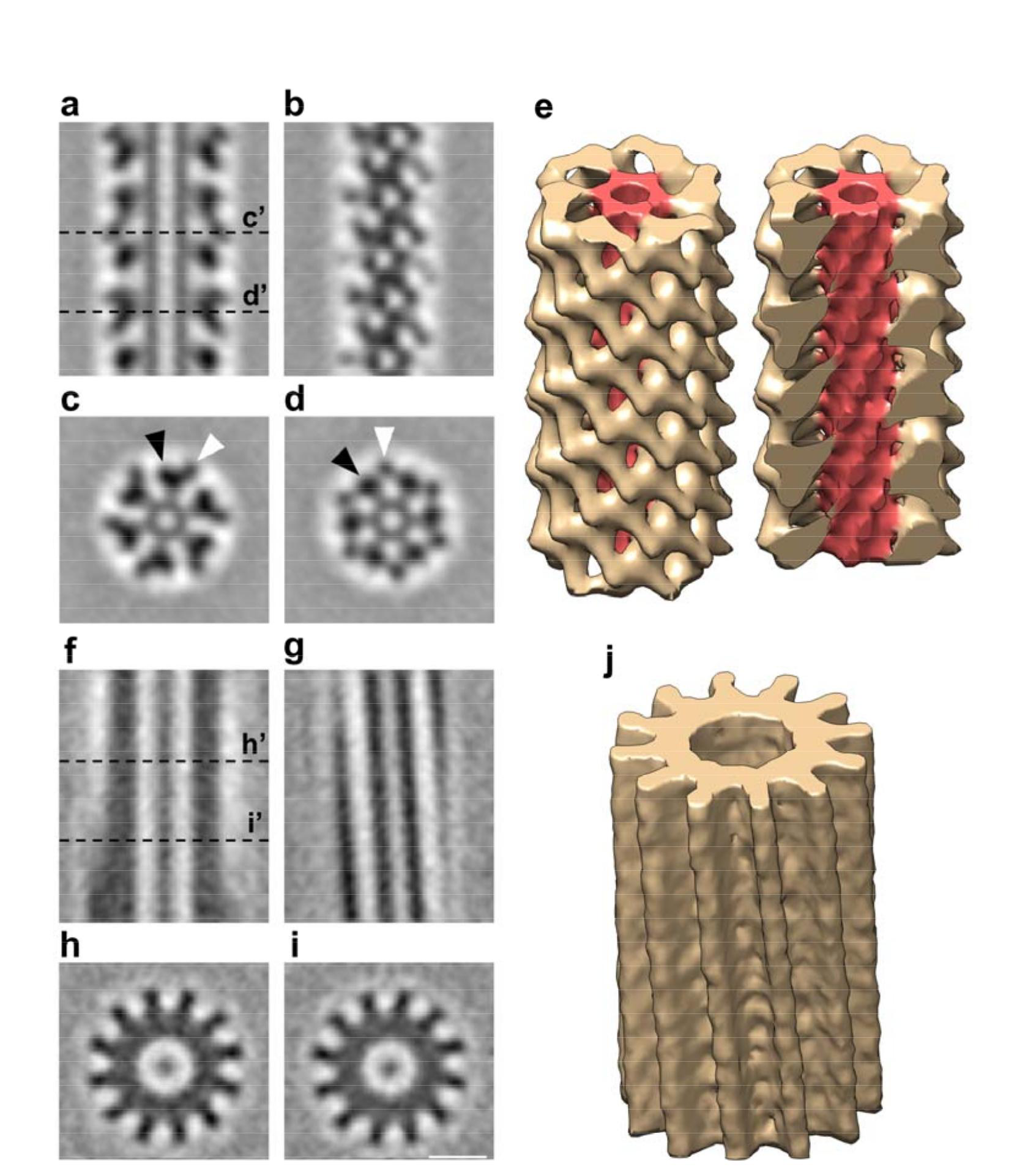
Sub-tomogram averages of the extended and contracted 381 T6SS sheaths in intact *M. xanthus* cells. (a, b) Slices through the center (a) and surface (b) of the sub-tomogram average of the extended T 6SS sheath. (c, d) Slices of cross-sections c’ and d’ of the sub-tomogram average shown in (a).Black and white arrowheads indicate the large and small domains of the sheath subunit, respectively. (e) 3-D envelope of the extended T 6SS sheath sub-tomogram average. The sheath is colored in yellow and the Hcp inner tube in red. Left: the full molecule is displayed; Right: a clipping plane is used to display the Hcp inner tube. (**f-j**) the same representations as in (a-e) for the contracted T6SS. Scale bar in (i) 10 nm, applies to (a-d, f-i).

Inspecting the more than 1,650 tomograms of *M. xanthus* cells in our tomography database^16^, we identified 29 extended T6SS with an average length of 444 nm (an example is shown in), and 24 contracted T6SS with an average length of 282 nm (an example is shown in Fig. 1b). Of the 29 extended T6SS structures, 16 were oriented close enough to horizontal (perpendicular to the electron beam) that we could align and average the membrane-associated region (Fig. 1c). The elongated structure of the sheath/Hcp tube was seen to end at a thin but distinct layer of density (L1). Another thicker layer of density (L2) was seen next to L1 with a gap in between. Between L2 and the IM, a cage-like structure (L3) was revealed, which surrounds a thin axial density with a bulge close to the IM. The L1, L2, and most of the L3 densities together likely represents the baseplate structure *in vivo*, and the thin axial density is likely the spike complex. A distinct density spanning the periplasm with a wider part close to the IM and a narrower part close to the OM was seen. This periplasmic density together with a part of the cytoplasmic L3 density likely represent the membrane complex, which associates with the OM and spans the periplasm and IM. An extracellular cap-like density was also seen on top of the OM outside of the cell

To date, no extracellular components of the T6SS have been identified. Therefore we were surprised to observe the ‘cap’. Moreover, in individual tomograms of T6SS we found densities resembling bacteriophage T4 tail fibers originating from the OM and extending into the extracellular environment (Fig. 1d). Of the 16 well-orientated extended T6SS, 13 exhibited tail fiber-like antennae. The number of antennae per T6SS varied from 1 to 6 with an average of 4. The bulb-like density on the tip of the antennae bears a strong resemblance to similar structures on the bacteriophage T4 tail fibers that contain the putative receptor interaction sites^17,18^. However, the average length of the antennae seen here is ~60 nm, different from the bacteriophage T4 long (145 nm) and short (34 nm) tail fibers^19^. At this point it is not clear if a part of the antennae spans the periplasm to connect with the baseplate, so the actual length of the antennae may be longer. There is no gene in the *M. xanthus* T6SS operon with sequence homology to bacteriophage tail fiber proteins, so the protein identity of these extracellular antennae remains unclear. Since the gene likely resides outside of the conserved T6SS operon, it could be a specialized accessory component that varies among different bacteria, conferring different target specificity and firing behaviors^2^.

To investigate the structure of the extended T6SS sheath, we computationally boxed the 29 extended sheaths observed in our cryotomograms into serial overlapping fragments and generated a sub-tomogram average. This revealed clear structural features of the sheath surrounding the Hcp inner tube (Fig 2a-d). The extended sheath exhibits six-fold symmetry along its long axis. In cross-sections of the sheath, each of the six lobes was seen to contain a large and a small domain (Fig. 2c, d; black arrowhead: large domain; white arrowhead: small domain). The large domains were connected to the inner tube by bridge densities; the small domains in turn connected to the large domains. A 3-D volume of the sub-tomogram average of the extended T6SS is shown in Fig. 2e. We determined a helical symmetry with a rise of 37Å and a rotation of 22 degrees for this six-start right-handed helix. We also used the same process to obtain a sub-tomogram average of 24 contracted T6SS sheaths observed in our cryotomograms (Fig. 2f-j). The cross-section of the contracted sheath showed a cogwheel-like structure with 12 distinct ridges (Fig. 2h, i), consistent with previous *in viv*o observations in *V. cholerae*

Sub-tomogram averages of the extended and contracted T6SS sheaths in intact *M. xanthus* cells.(a, b) Slices through the center (a) and surface (b) of the sub-tomogram average of the extended T6SS sheath. (c, d) Slices of cross-sections c’ and d’ of the sub-tomogram average shown in (a). Black and white arrowheads indicate the large and small domains of the sheath subunit, respectively. (e) 3-D envelope of the extended T6SS sheath sub-tomogram average. The sheath is colored in yellow and the Hcp inner tube in red. Left: the full molecule is displayed; Right: a clipping plane is used to display the Hcp inner tube. (f-j) the same representations as in (a-e) for the contracted T6SS. Scale bar in (i) 10 nm, applies to (a-d, f-i).

Since it is unclear how a higher resolution structure of the extended T6SS sheath might be obtained directly, and since a wealth of structural information is available for the R-type pyocin and contracted T6SS, we generated a pseudo-atomic model of the extended T6SS sheath by applying this known information to our sub-tomogram average. We started by superposing an atomic model of the extended R-type pyocin (PDB 3J9Q) onto the sub-tomogram average of the extended sheath using the Hcp inner tube as a reference (Fig. 3a). In the extended R-type pyocin structure, the sheath protein interacts with Hcp through an attachment α-helix close to the C-terminus^11^, which is also present in the *M. xanthus* T6SS sheath protein TssC. In our sub-tomogram average we see a clear connection between the sheath and Hcp densities, which is well occupied by the attachment α-helix (Fig. 3a, arrow). The other parts of the pyocin sheath protein structure do not fit the density well, since the T6SS sheath is formed not by a single protein but by a TssB-TssC heterodimer and contains an extra recycling domain. Near-atomic-resolution structures of the TssB-TssC heterodimer were resolved previously in contracted T6SS sheaths purified from *V. cholerae* and *Francisella novicida*^12,13^. We used the more closely related *V.cholerae* structure (PDB 3J9G) as a template to generate a homology model of the *M. xanthus* TssB-TssC heterodimer (MxTssBC) with which to replace the pyocin sheath protein structure. We superposed the MxTssBC model onto the pyocin sheath protein based on their structurally conserved domains^11^ (Fig. 3b). The result of this superposition suggests that the T6SS TssB-TssC likely uses the same attachment α-helix to interact with the Hcp protein in the inner tube (Fig. 3b, arrow), but the MxTssBC model did not fit the density best in this orientation. It has been shown that upon pyocin sheath contraction, the sheath subunits rotate as rigid bodies. We therefore hypothesized that the structure of individual MxTssBC subunits remains the same in the extended sheath as in the contracted one. We then treated MxTssBC as a rigid body and slightly rotated it to best fit the density while maintaining the location of the attachmentα-helix for interaction with Hcp (Fig. 3c, d). We also replaced the Hcp structure with a homology model of the *M. xanthus* Hcp hexamer (MxHcp) (constructed using PDB 3EAA^20^ as a template) (Fig. 3d). The conserved structural features at the interface of MxTssBC and MxHcp suggest that the sheath and Hcp tube in the T6SS likely interact in the same manner as in the R-type pyocin (Fig. 3a, d). We then replicated the MxTssBC model to populate the remaining five MxHcp subunits to create a hexamer of MxTssBC-MxHcp, which constitutes one layer of the extended T6SS sheath (Fig. 3d).

**Figure 3.**
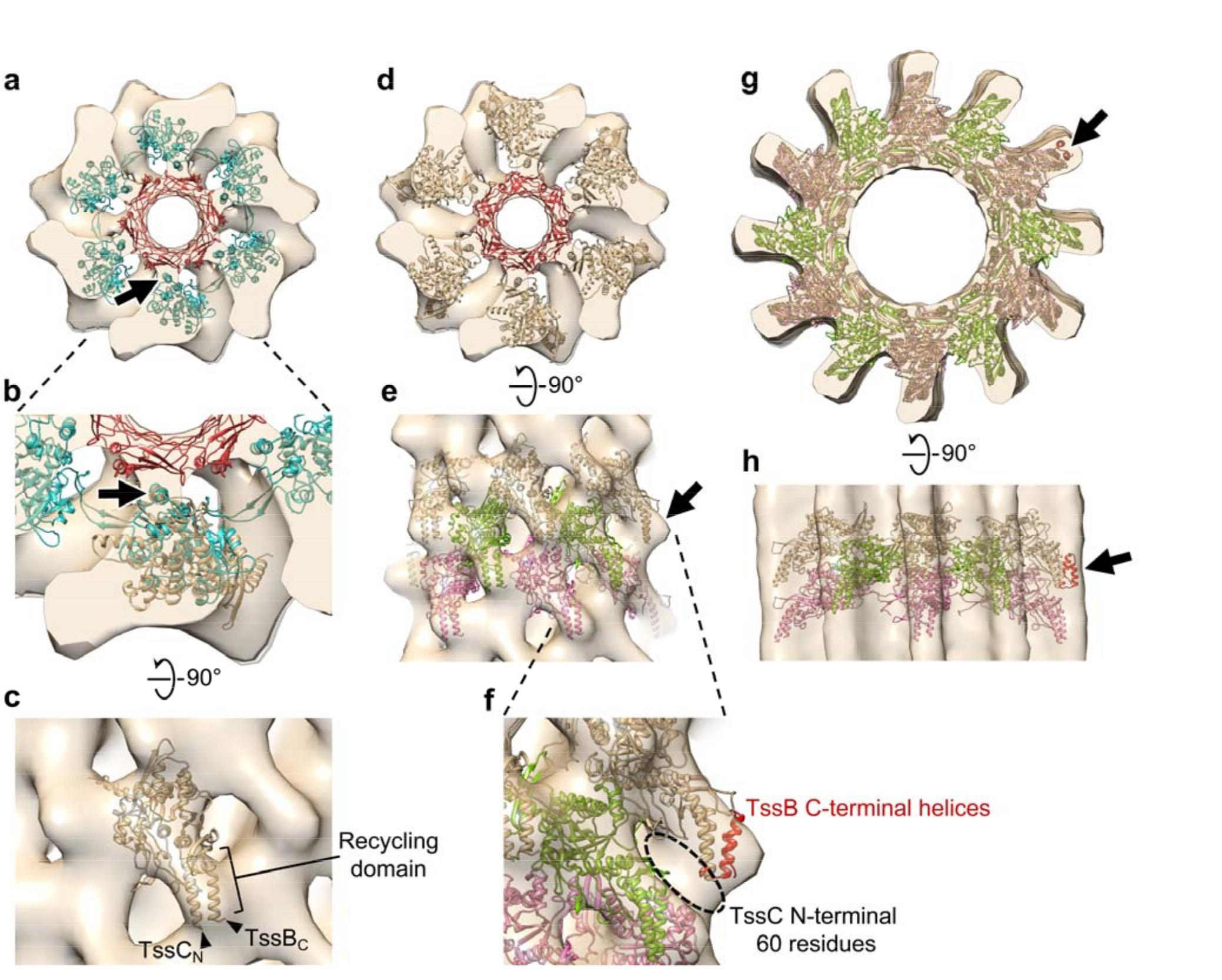
Generating pseudo-atomic models of the *M. xanthus* T6SS sheath in extended and contracted conformations. (a) An atomic model of the extended R-type pyocin (PDB 3J9Q) placed in the envelope of the extended T6SS sheath sub-tomogram average, fitting the Hcp inner tube (red) and the attachmentα-helix (arrow) in the sheath (cyan) to corresponding densities. The envelope shown here contains two layers of the extended sheath. (b) Enlarged view of part of (a), with an MxTssBC model superposed on one sheath subunit of the R-type pyocin structure. Arrow: the conserved attachment α-helix in T6SS and R-type pyocin sheath subunits. (c) Side view of (b) after the MxTssBC model was rotated to best fit the density. TssCN and TssBC indicate the N-terminus of and C-terminus of TssB, respectively. (d) The model of h 400 exameric MxTssBC-MxHcp (one layer of the extended T6SS sheath) placed in the envelope of the extended M. xanthus T6SS sheath sub-tomogram average. The envelope shown here contains two layers of the extended sheath. (e) Side view of (d) after fitting three consecutive layers of the hexameric MxTssBC404 p model (shown in different colors) into the envelope. Arrow: density not fully occupied by the model. (f) Enlarged view of part of (e), with the model of TssB C-terminal helices (red; generated using PDB 4PS2) concatenated to the model in (e). Dashed circle: densities likely occupied by the 60 N-terminal TssC amino acid residues, whose structure was absent in the MxTssBC model. (g) The homology model of the contracted M. xanthus T6SS sheath (generated using PDB 3J9G) placed into the envelope of the contracted T6SS sheath sub-tomogram average. (h) Side view of (g), with three consecutive layers of hexameric MxTssBC-MxHcp shown in different colors. Arrows in (g) and (h) indicate additional densities, which can be partially filled by concatenating the model of TssB C-terminal helices (red) as in (f). Note that both red helices are visible in (g) and (h), but due to view angle, they appear as one in (f).

Next we propagated the MxTssBC-MxHcp hexamer down the length of the sheath with a 37Å translation and 22 degree rotation of each new layer to fit the density (Fig. 3e). Most of the density was occupied, with the exception of the region surrounding the TssC N-terminus and TssB C-terminus in the MxTssBC model (Fig. 3e, arrow). This structure comprises the recycling domain of the T6SS sheath (Fig. 3c). A portion of this recycling domain was not resolved in the high-resolution (<3.5Å) structures of contracted T6SS sheaths^12,13^ that we used to generate the MxTssBC model. An earlier lower-resolution (6Å) cryo-EM reconstruction of the *V. cholerae* T6SS contracted sheath revealed that this missing part is composed of at least two helical elements and some additional densities^21^. A crystal structure of the TssB C-terminal domain is available (PDB 4PS2) that partially overlaps with the TssB C-terminus in the MxTssBC model and contains the remaining two helices of the recycling domain. We concatenated this structure with the MxTssBC model and observed that it occupies the additional density well (Fig. 3f). The additional ~60 residues of the TssC N-terminus absent in the MxTssBC model likely occupy the remaining empty densities (Fig. 3f, dashed circle).

We next constructed a pseudo-atomic model of the contracted sheath by placing a model of MxTssBC, generated based on the packing observed in the *V. cholerae* contracted sheath structure (PDB 3J9G)^12^, into our sub-tomogram average. The inner diameter and dodecameric ridges of the model fit the density well (Fig. 3g, h). Interestingly, we observed additional densities at the tips of the ridges. As mentioned above, these densities were not observed in high-resolution reconstructions of purified contracted sheaths^12,13^, but were seen in a lower-resolution reconstruction^21^. This suggests that the tip regions might exhibit greater flexibility than other parts. In our sub-tomogram average, these additional densities could be partially occupied by the TssB C-terminal helices (Fig. 3g, h, arrows), as we saw for the extended conformation.

The pseudo-atomic models of extended and contracted *M. xanthus* T6SS sheaths generated here support the hypothesis that the T6SS and R-type pyocin share a similar architecture and mechanism (Fig. 4). In our models the protofilaments of the T6SS undergo similar motions to pyocin upon sheath contraction (Fig. 4b, e, h, k), but when the same number of layers of the T6SS and pyocin are compared, the pyocin shows greater contraction (~33% shortening in the T6SS vs. ~56% in pyocin). However, since the length of extended T6SS can be more than 4X the length of yocins^11^, T6SS can still penetrate deeper into the target cell.

**Figure 4.**
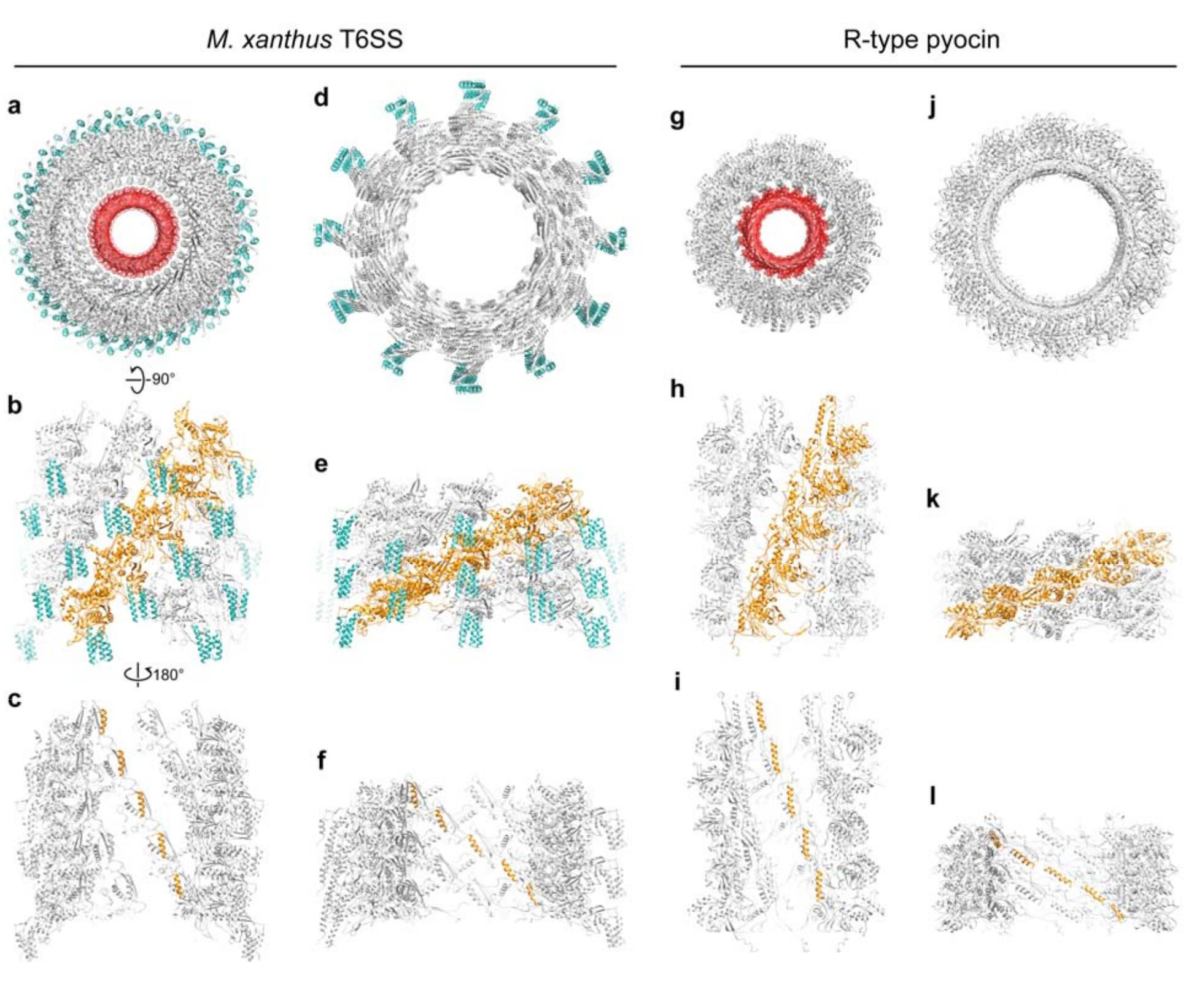
Comparing the models of *M. xanthus* T6SS sheath 415 and R-type pyocin in different conformations. (a) Top view of the extended M. xanthus T6SS sheath model containing 5 layers of MxTssBC418 MxHcp hexamers. The sheath is colored gray, with the recycling domain colored cyan. The Hcp inner tube is colored red. (b) Side view of the model in (a), with only the front half shown for clarity. One protofilament of the sheath is colored orange. (c) Back view (from inside the tube) of model shown in (b). Here only the attachment α-helices in one protofilament of the sheath are colored orange. (d-f) The same representations as in (a-c) but for the contracted M. xanthus T6SS sheath. (g-l) The same representations as in (a-f) but for the extended R-type pyocin.

The extended pyocin structure reported in a previous study revealed details of the interactions between the sheath and Hcp tube, and showed that the Hcp tube packs in a helical fashion like the sheath^11^. In the T6SS, however, how the Hcp hexamers stack in the inner tube is still indebate^7^. Although our sub-tomogram average of the extended T6SS is of insufficient resolution to reveal clear features of the Hcp hexamers in the inner tube, the helical arrangement of the density for the putative Hcp-interacting α-helix on the inside of the T6SS sheath reveals that Hcp hexamers in the T6SS are also packed helically with the symmetry of the sheath (Fig. 4c, f, i, l).

While many aspects of the T6SS structure and contraction mechanism are clearly conserved with the R-type pyocin, a key difference is the presence of recycling domains on the T6SS sheath (Fig. 4a, b, d, e). As discussed above, in the contracted T6SS sheath, the recycling domain is exposed at the tip of the sheath ridges and is less conformationally rigid, possibly allowing access to the ATPase ClpV for sheath disassembly. Here we show that in the extended sheath, the recycling domain is partly obstructed by interaction with the neighboring protofilament (Fig. 4b). We speculate that this protection of the recycling domain prevents premature disassembly of the extended sheath by ClpV. In addition, in the extended pyocin structure, most of the interactions among sheath subunits are confined to within a single protofilament. Only the extended flexible arms on the N-and C-termini of sheath subunits link neighboring protofilaments^11^ (Fig. 4h). Here we observed that the recycling domain of the T6SS sheath is sandwiched in between two neighboring protofilaments (Fig. 4b), suggesting that this domain might also be involved in assembly and stabilization of the extended sheath.

Finally, we placed our new sub-tomogram average and the previously solved structures of individual components into the overall density (Fig. 5a-c). The baseplate has been proposed to consist of TssA, TssE, TssF, TssG and TssK^1^. While TssA is initially recruited near the membrane, it has recently been shown to remain at the growing end (away from the membrane) of extending sheaths as new sheath and Hcp subunits are incorporated^22^, and so is likely not present in the mature baseplate. It has been proposed that the T6SS sheath associates with the baseplate through TssE, because TssE is required for TssBC assembly^4^ and shows clear homology to gp25 in the bacteriophage baseplate, which is known to interact with the bacteriophage sheath^23^. Since TssE only comprises 131 residues and was proposed to interact with the sheath through a continuation of the three-way ‘handshake’ interactions of subunit domains in the assembled sheath^12^, we reasoned that the density of TssE would be small and directly connected to the sheath density without any gap, similar to the gp25 location in the reported bacteriophage structure^23^. We therefore proposed that the small but distinct density layer L1 is TssE and placed the average of the extended sheath/Hcp tube abutting it (Fig. 5c). It is also possible that a part of the L1 density is contributed by another baseplate component TssK, which has been shown to directly interact with Hcp, TssC of the sheath and TssL of the membrane complex, and therefore it was proposed to connect the sheath/inner tube to the membrane complex^24^. Purified TssK was previously shown to form a trimeric structure insolution with dimensions of around 100 × 100 × 90 Å^24^. Based on our sub-tomogram average, in order to span through the L1, L2 and L3 density layers (~300 Å) to interact with both the membrane complex and the sheath/Hcp tube, TssK must exhibit a more extended conformation or form a higher-order oligomer in the assembled baseplate *in vivo* than in the isolated structure *in vitro*

**Figure 5.**
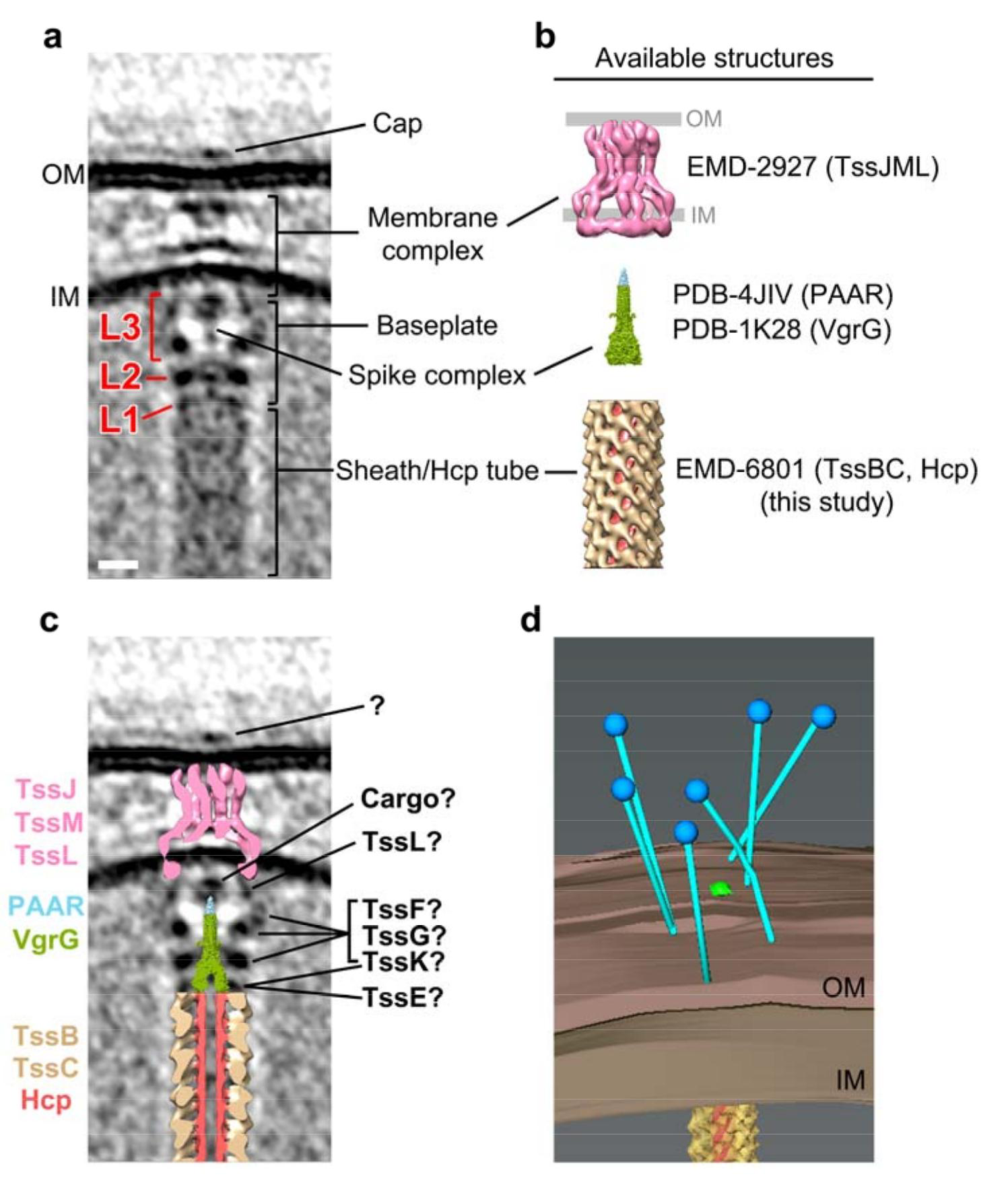
Placing available component structures into 426 the *M. xanthus* T6SS structure *in vivo*. (a) Central slice of the subtomogram average of M. xanthus T6SS membrane-associated region. (b) Available structures and models of individual components. The membrane complex structure (EMD-2927) is shown with the OM and IM locations defined by previous work29. The spike complex model was generated by concatenating the structure of VgrG from bacteriophage T4 (PDB 1K28) with non-homologous parts removed with the crystal structure of the T6SS PAAR433 repeat protein (PDB 4JIV). The extended sheath/Hcp tube structure was generated in this study (EMD-8601). (c) Placing the central slices of available structures 434 into the average shown in (a). (d) 3-D segmentation of the bacteriophage tail fiber-like antennae shown in Fig.1d (blue:bacteriophage tail fiber-like antennae; brown: inner and outer membranes; yellow: sheath; red:Hcp inner tube). Scale bar in (a) 10 nm, applies to (a) and (c).

We next generated a homology model of the spike complex by concatenating the structure of VgrG from bacteriophage T4 (PDB 1K28)^25^ with the crystal structure of the T6SS PAAR-repeat protein (PDB 4JIV)^26^ (Fig. 5b). VgrG was shown to interact directly with the Hcp tube^27^. We therefore placed the model of the spike complex adjacent to the tip of the fitted Hcp tube (Fig. 5c). The model fits its potential density well in the center of baseplate, but interestingly the tip portion containing the PAAR-repeat protein is clearly smaller than its corresponding density in the sub-tomogram average. It has been proposed that some T6SS cargo effectors may associate with the PAAR-repeat protein for delivery^2,26,28^. We therefore speculate that cargo effectors may partly contribute to this density (Fig. 5c). Recently, the T6SS membrane complex, composed of TssJ and TssM in the periplasm and TssL mainly in the cytoplasm, was purified and visualized by single particle cryo-EM^29^. The overall shape and dimensions of this single particle structure fit our *in vivo* densities well. However, the potential TssL density in the cytoplasm is more extended in our sub-tomogram average, perhaps due to better structural preservation in the native environment and intact association with baseplate proteins (Fig. 5c). The remaining unassigned baseplate densities in L2 and L3 are likely contributed by the baseplate proteins TssF, TssG and TssK (Fig. 5c). The extracellular bacteriophage tail fiber-like antennae which seen in individual T6SS (Fig. 1d) is shown by a 3-D segmentation in Fig. 5d.

In summary, here we have presented the first sub-tomogram average of an extended T6SS sheath structure, revealing new insights into how its recycling domains are regulated. We also present the first visualization of the baseplate structure, revealing how it associates with the membrane complex and anchors the sheath. This structure also provides evidence of how cargo effectors might associate with the PAAR-repeat protein of the spike complex. Our observation of extracellular antennae opens a new path for future research into how this important nanomachine recognizes targets and triggers firing. In the future, it will also be of great interest to locate all the components in the T6SS structure by imaging mutants (for instance as we did for the type IV pilus machines^30,31^). *M. xanthus* is not a good system for such mutant analysis, however, because of the low number of T6SS in the cells (we only obtained 16 T6SS structures showing clear views of the membrane-associated components from >1,650 tomograms) and the large size of the cells. New model systems such as T6SS-containing minicells should therefore be sought for further dissection and higher-resolution analysis of the T6SS *in vivo*

## METHODS

### Electron cryotomography

*M. xanthus* DK 1622 strain was grown in 10 mL CTT medium at 32°C with 250 rpm shaking. The cells were mixed with 10-nm colloidal gold (Sigma-Aldrich, St. Louis, MO) pretreated with bovine serum albumin and subsequently applied to freshly glow-discharged Quantifoil copper R2/2 200 EM grids (Quantifoil Micro Tools GmbH, Jena, Germany). The grids were plunge-frozen in a liquid ethane propane mixture^32^ using an FEI Vitrobot Mark III (Thermo Fisher Scientific, Waltham, MA). The frozen grids were imaged in an FEI Polara 300 keV field emission gun transmission electron microscope (Thermo Fisher Scientific, Waltham, MA) equipped with a Gatan energy filter (Gatan, Pleasanton, CA) and a Gatan K2 Summit direct detector (Gatan, Pleasanton, CA). Energy-filtered tilt-series of images were collected automatically from −60° to +60° at 1° intervals using the UCSF Tomography data collection software^33^ with a total dosage of 160 e^-^/Å^2^, a defocus of −6 µm and a pixel size of 3.9 Å. The images were aligned and contrast transfer function corrected using the IMOD software package^34^. SIRT reconstructions were then produced using the TOMO3D program^35^. T6SS structures were located by visual inspection. The sub-tomogram averages were produced using

The PEET program^36^. To generate the sub-tomogram averages of the T6SS sheaths, model points were distributed along the sheaths for the PEET program to crop out and align the overlapping boxes of densities.

## Data availability

The sub-tomogram averages of the *M. xanthus* T6SS that support the findings of this study have been deposited in the Electron Microscopy Data Bank with accession codes EMD-8600 (membrane-associated region); EMD-8601 (extended sheath); EMD-8602 (contracted sheath). The coordinates of the extended and contracted sheath models have been deposited in the Protein Data Bank with accession code 5URW and 5URX, respectively.

## ACKNOWLEDGEMENTS

We thank Prof. Hong Z. Zhou and Dr. Peng Ge for providing the initial concept of how to place the MyTssBC model in the sub-tomogram average of extended T6SS sheath and Dr. Catherine Oikonomou for discussions and editorial assistance. This work was supported by NIH grant R01 AI127401 to G. J. J.

## CONTRIBUTIONS

Y.-W. C. and L. A. R. collected, processed and analyzed the electron cryotomography data. Y.-W. C. built the final models. Y.-W. C. and G. J. J. wrote the paper.

---

**Figure S1.**
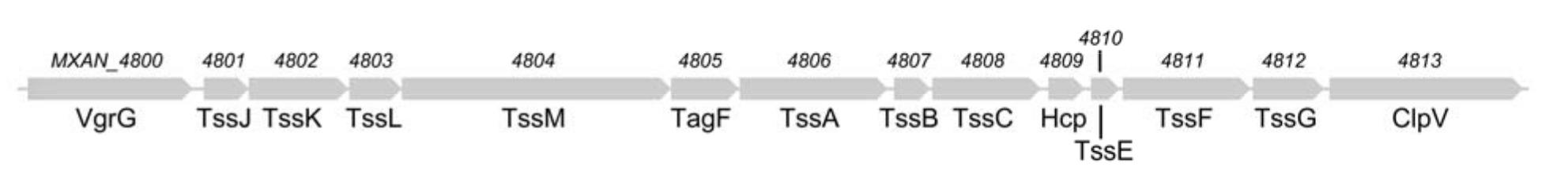
M. xanthus T6SS gene cluster. The cluster of conserved T6SS genes (MXAN_4800 to MXAN_4813) in the genome of M.xanthus.

**Figure S2.**
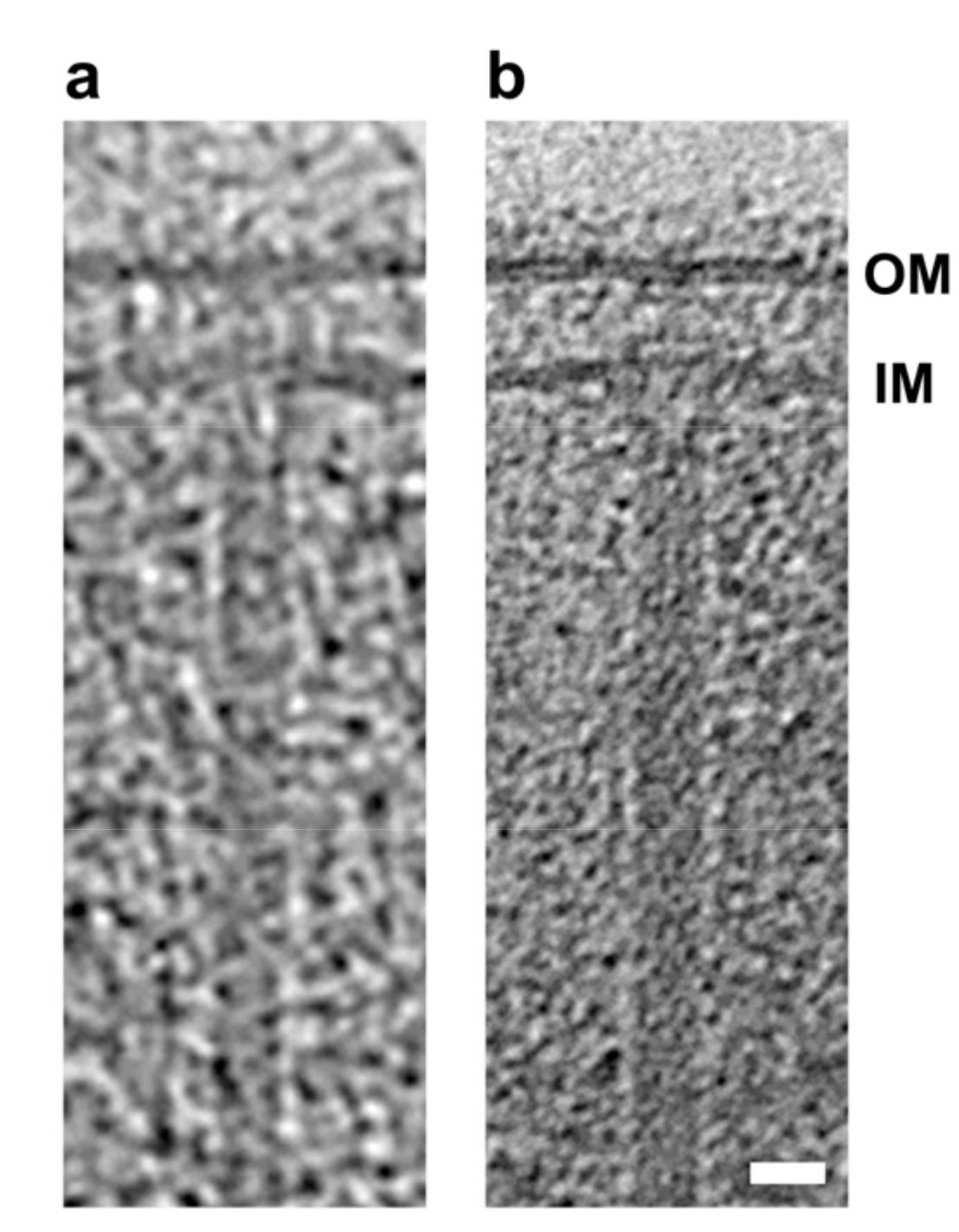
Images of M. xanthus 443 T6SS at different defoci. (a) A T6SS structure captured by correlated cryo-PALM/ECT in our previous study15 using a defocus of -10 µm during data collection. (b) A similar structure captured in a cryotomogram of an M. xanthus cell with a defocus of −6 µm. Tomography slices through full 3-D reconstructions of intact cells are shown in both cases. Scale bar in (b) 20 nm, applies to (a) and (b).

## REFERENCES

1 Cianfanelli,F. R., Monlezun,L. & Coulthurst,S. J. Aim, Load,Fire: The Type VI Secretion System, a Bacterial Nanoweapon. Trends in microbiology 24, 51–62 (2016).

2 Ho, B. T., Dong, T. G. & Mekalanos, J. J. A view to a kill: the bacterial type VI secretion system. Cell host & microbe 15, 9–21 (2014).

3 Zoued, A. et al. Architecture and assembly of the Type VI secretion system. Biochimica et biophysica acta 1843, 1664–1673 (2014).

4 Basler, M., Pilhofer, M., Henderson, G. P., Jensen, G. J. & Mekalanos, J. J. Type VI secretion requires a dynamic contractile phage tail-like structure. Nature 483, 182–186 (2012).

5 Basler, M., Ho, B. T. & Mekalanos, J. J. Tit-for-tat: type VI secretion system counterattack during bacterial cell-cell interactions. Cell 152, 884–894 (2013).

6 Basler, M. & Mekalanos, J. J. Type 6 secretion dynamics within and between bacterial cells. Science 337, 815 (2012).

7 Kube, S. & Wendler, P. Structural comparison of contractile nanomachines. AIMS Biophysics, 88-115 (2015).

8 Vettiger, A. & Basler, M. Type VI Secretion System Substrates Are Transferred and Reused among Sister Cells. Cell 167, 99–110, e112 (2016).

9 Pietrosiuk, A. et al. Molecular basis for the unique role of the AAA+ chaperone ClpV in type VI protein secretion. The Journal of biological chemistry 286, 30010–30021 (2011).

10 Kapitein, N. et al. ClpV recycles VipA/VipB tubules and prevents non-productive tubule formation to ensure efficient type VI protein secretion. Molecular microbiology 87, 1013–1028 (2013).

11 Ge, P.et al. Atomic structures of a bactericidal contractile nanotube in its pre-and postcontraction states. Nature structural & molecular biology 22, 377–382 (2015).

12 Kudryashev, M. et al. Structure of the type VI secretion system contractile sheath. Cell 160, 952–962 (2015).

13 Clemens, D. L., Ge, P., Lee, B. Y., Horwitz, M. A. & Zhou, Z. H. Atomic structure of T6SS reveals interlaced array essential to function. Cell 160, 940–951 (2015).

14 Oikonomou, C. M., Chang, Y.-W. & Jensen, G. J. A new view into prokaryotic cell biology from electron cryotomography. Nature reviews. Microbiology 14, 205–220 (2016).

15 Chang, Y. W. et al. Correlated cryogenic photoactivated localization microscopy and cryo-electron tomography. Nature methods 11, 737–739 (2014).

16 Ding, H. J., Oikonomou, C. M. & Jensen, G. J. The Caltech Tomography Database and Automatic Processing Pipeline. Journal of structural biology 192, 279–286 (2015).

17 Bartual, S. G. et al. Structure of the bacteriophage T4 long tail fiber receptor-binding tip. Proceedings of the National Academy of Sciences of the United States of America 107, 20287–20292 (2010).

18 Thomassen, E. et al. The structure of the receptor-binding domain of the bacteriophage T4 short tail fibre reveals a knitted trimeric metal-binding fold. Journal of molecular biology 331, 361–373 (2003).

19 Yap, M. L. & Rossmann, M. G. Structure and function of bacteriophage T4. Future microbiology 9, 1319–1327 (2014).

20 Jobichen, C. et al. Structural basis for the secretion of EvpC: a key type VI secretion system protein from Edwardsiella tarda. PloS one 5, e12910 (2010).

21 Kube, S. et al. Structure of the VipA/B type VI secretion complex suggests a contraction-state-specific recycling mechanism. Cell reports 8, 20–30 (2014).

22 Zoued, A. et al. Priming and polymerization of a bacterial contractile tail structure. Nature 531, 59–63 (2016).

23 Taylor, N. M. et al. Structure of the T4 baseplate and its function in triggering sheath contraction. Nature 533, 346–352 (2016).

24 Zoued, A. et al. TssK is a trimeric cytoplasmic protein interacting with components of both phage-like and membrane anchoring complexes of the type VI secretion system. The Journal of biological chemistry 288, 27031–27041 (2013).

25 Kanamaru, S. et al. Structure of the cell-puncturing device of bacteriophage T4. Nature 415, 553–557 (2002).

26 Shneider, M. M. et al. PAAR-repeat proteins sharpen and diversify the type VI secretion system spike. Nature 500, 350–353 (2013).

27 Lin, J. S., Ma, L. S. & Lai, E. M. Systematic dissection of the agrobacterium type VI secretion system reveals machinery and secreted components for subcomplex formation. PloS one 8, e67647 (2013).

28 Koskiniemi, S. et al. Rhs proteins from diverse bacteria mediate intercellular competition. Proceedings of the National Academy of Sciences of the United States of America 110, 7032–7037 (2013).

29 Durand, E. et al. Biogenesis and structure of a type VI secretion membrane core complex. Nature 523, 555–560 (2015).

30 Chang, Y. W. et al. Architecture of the type IVa pilus machine. Science 351, aad2001 (2016).

31 Chang, Y. W. et al. Architecture of the Vibrio cholerae toxin-coregulated pilus machine revealed by electron cryotomography. Nature microbiology 2, 16269 (2017).

32 Tivol, W. F., Briegel, A. & Jensen, G. J. An improved cryogen for plunge freezing. Microscopy and microanalysis: the official journal of Microscopy Society of America, Microbeam Analysis Society, Microscopical Society of Canada 14, 375–379 (2008).

33 Zheng, S. Q. et al. UCSF tomography: an integrated software suite for real-time electron microscopic tomographic data collection, alignment, and reconstruction. Journal of structural biology 157, 138–147 (2007).

34 Kremer, J. R., Mastronarde, D. N. & McIntosh, J. R. Computer visualization of three-dimensional image data using IMOD. Journal of structural biology 116, 71–76 (1996).

35 Agulleiro, J. I. & Fernandez, J. J. Fast tomographic reconstruction on multicore computers. Bioinformatics 27, 582–583 (2011).

36 Nicastro, D. et al. The molecular architecture of axonemes revealed by cryoelectron tomography. Science 313, 944–948 (2006).

